# Live-tracking of replisomes reveals nutrient-dependent regulation of replication elongation rates in *Caulobacter crescentus*

**DOI:** 10.1101/2024.08.20.608768

**Authors:** Inchara S Adhikashreni, Asha Mary Joseph, Sneha Phadke, Anjana Badrinarayanan

**Affiliations:** National Centre for Biological Sciences (TIFR), Bengaluru 560065, India

**Keywords:** DNA replication, replisome dynamics, *Caulobacter crescentus*, live-cell imaging, stationary phase, replication regulation

## Abstract

In bacteria, commitment to genome replication (initiation) is intricately linked to nutrient availability. Whether growth conditions affect other stages of replication beyond initiation, remains to be systematically studied. To address this, we assess the replication dynamics of *Caulobacter crescentus*, a bacterium that undergoes only single round of replication per cell cycle, by tracking the replisome across various growth phases and nutrient conditions. We find that the replication elongation rates slowdown as cells transition from exponential (high-nutrient) to stationary (low-nutrient) phase, and this contributes significantly to the overall cell cycle delay. While elongation rates are correlated with growth rates, both properties appear to be differentially influenced by nutrient status. This slowdown in replication progression is not associated with increased mutagenesis or upregulation of the DNA damage responses. We conclude that the growth conditions not only dictate the commitment to replication but also the rates of genome duplication. Such regulation appears to be distinct from stress-induced replication slowdown and likely serves as an adaptive mechanism to cope with fluctuations in nutrient availability in the environment.

## Introduction

Chromosome replication is essential for the propagation of life, and is carried out by a multi-subunit complex called the replisome (Xu & Dixon, 2018; Yao & O’Donnell, 2016). In bacteria, replication initiates at the origin of replication (*‘ori’*). Two replisomes carry out DNA synthesis bidirectionally along the chromosome arms and replication terminates when the replisomes meet at the terminus (*‘ter’*) (Bates & Kleckner, 2005; Berkmen & Grossman, 2006; Gyurasits & Wake, 1973; Jensen, 2001; Prescott & Kuempel, 1972; Reyes-Lamothe et al., 2008; Zhang et al., 2024). These steps of replication can be used to define the bacterial cell cycle stages: a. Initiation (B period) entails the time from cell birth until the start of DNA replication. b. Elongation (C period) is the period of DNA synthesis and c. Division (D period) is the time between completion of replication and cell division (Skarstad et al., 1983). DNA replication must be integrated with cell growth and environmental fluctuations, to ensure the transmission of a fully replicated copy of the genome at the time of cell division. While this appears to be a challenging task given large variations in growth rates and/ or nutrient availability, replication is remarkably robust and accurate (Govers et al., 2023). How bacteria maintain control and coordination of DNA replication and cell cycle progression across diverse growth conditions is still not completely understood.

Although many bacteria lack dedicated cell cycle-checkpoint mechanisms (as observed in eukaryotes), the commitment to replication (initiation) is highly regulated via multiple mechanisms (Frandi & Collier, 2019; Hallgren & Jonas, 2024; Skarstad & Katayama, 2013). For example, in fast growing bacteria such as *E. coli*, which divide faster than their C period in nutrient rich conditions, new rounds of replication are initiated in previous cell cycle itself (Cooper & Helmstetter, 1968). Control of initiation is centered on the replication initiation protein DnaA, and this has been extensively studied, both during steady state growth in different nutrient conditions as well as in contexts of stress such as starvation and stationary phases (Collier & Shapiro, 2009; Felletti et al., 2021; Hallgren et al., 2023; Jonas et al., 2013; Kato & Katayama, 2001; Leslie et al., 2015; Liu et al., 2016; Noirot-Gros et al., 2006). Apart from initiation, it is unclear whether other stages of replication are also regulated based on nutrient availability.

Our understanding of replication control has originally been derived from population-level experiments, from which also emerged a systems-level understanding of the intricate relationship between cell size, growth rate and cell cycle regulation (Cooper & Helmstetter, 1968; Donachie, 1968; Schaechter et al., 1958). However, precise measurement of elongation rates across diverse growth phases/ conditions has been challenging. While earlier studies estimating C period at the population-level in *E. coli* proposed invariance across varied nutrient conditions, later ones suggested that this applied to only fast growth conditions, with both C and D periods also increasing as growth rates decreased (Allman et al., 1991; Bipatnath et al., 1998; Cooper & Helmstetter, 1968; Helmstetter & Cooper, 1968; Helmstetter & Pierucci, 1976; Michelsen et al., 2003; Sanders et al., 2023; Skarstad et al., 1983; Stokke et al., 2012). Paradoxically most studies that have probed replication timing are also limited to bacteria like *E. coli* and *Bacillus subtilis* which pose an additional challenge, as these systems, depending on growth conditions, initiate single or multiple rounds of replication within one cell cycle. Such ensemble measurements also mask cell-to-cell variations within the population, which have become evident with more recent single cell analysis. Apart from recapitulating growth and size laws at the level of single cells, these studies have provided valuable mechanistic insights into regulatory principles and have identified deviations from idealized population averages (Adiciptaningrum et al., 2015; Govers et al., 2023; Sanders et al., 2023; Si et al., 2019; Wallden et al., 2016). Incidentally the focus of these experiments has also been restricted to initiation control, or to broadly infer an overall cell cycle timing. It is possible that direct visualization of individual replication cycles and comparison of C periods across nutrient conditions becomes difficult in bacteria that can go through multiple initiation events within the same cell cycle. Indeed, it would be surprising if growth conditions do not impact other stages of replication, as this would imply that elongation would need to progress (at the same rates) to completion regardless of the growth context.

Here, we used *Caulobacter crescentus*, a bacterium that undergoes only a single round of replication in every cell cycle (Marczynski, 1999), to precisely and directly measure the replication timings. Using single-cell fluorescence time-lapse microscopy, we track the replisome across changing growth phases (from exponential to stationary) and nutrient conditions (rich vs. minimal media). We find that the cell cycle period increases as cells transition from exponential (high-nutrient) to stationary phase (low-nutrient) growth. Apart from shut-down of replication initiation, replication elongation times also increase, significantly contributing to the overall cell cycle delay. We observe a strong correlation between replication elongation rates and cellular growth rates, with both rates changing differentially in response to nutrient availability. Consistently, shifting cells from low-nutrient to high-nutrient conditions results in faster progression of replication elongation and increased cellular growth rates. Such nutrient-dependent slowdown in replication progression as cells enter stationary phase is not due to increased mutagenesis or upregulation of the DNA damage response, typically seen much later in stationary phase. Together, our study provides the first direct measure of replication progression as cells transition from exponential to stationary phase. Using this paradigm, we uncover a relationship between replication speeds and nutrient availability, highlighting that growth environment not only governs the commitment to replication but also the rates of genome duplication.

## Results

### Measurement of *Caulobacter* cell cycle stages via single-cell tracking of replisome dynamics

To track replication dynamics in real-time, we imaged the β-clamp subunit of the replisome (DnaN) fused to YFP, expressed from its endogenous locus. This strategy has been used previously to track replisome mobility inside single cells (Joseph et al., 2021; Mangiameli et al., 2017; Moolman et al., 2014; Reyes-Lamothe et al., 2008; Si et al., 2019; Zhang et al., 2024). As reported earlier, we observed no localizations of DnaN (diffuse cytoplasmic signal) in new-born *Caulobacter* cells (Jensen, 2001; Zhang et al., 2024). Following this, a single localization of DnaN was observed at a cell pole (where the *ori* is located), indicative of replication initiation. As replication progressed, the DnaN localization moved towards the opposite pole, and on completion of replication, dissociated around mid-cell where the *ter* is located. Such dissociation of DnaN localization is indicative of replication completion, and is followed by successful cell division (Fig. 1A, S1A-C). Though replication is mediated simultaneously by two replisomes on both chromosome arms, due to chromosome inter-arm alignment, only a single localization was visualized during most part of the replication cycle (Zhang et al., 2024). Since only a single round of replication is initiated in each cell cycle, by tracking the formation, progression and dissociation of the replisome localization, we can accurately measure each stage of replication – the time to form a fluorescent DnaN localization after cell birth (B period), the time period for which DnaN remains localized and is mobile in the cell (C period), and the time between dissociation of the DnaN localization and cell division (D period). We define ‘interdivision’ time or ‘cell cycle’ time as the total time taken between two cell divisions (B+C+D periods) (Fig. 1A, S1A-C). For example, in exponentially growing cells (OD_600_ 0.2-0.4), B period of 4.6 ± 4 min, C period of 69.8 ± 7.4 min and D period of 5.8 ± 4.6 min correspond to an interdivision time of 80.2 ± 8.8 min.

**Figure 1:**
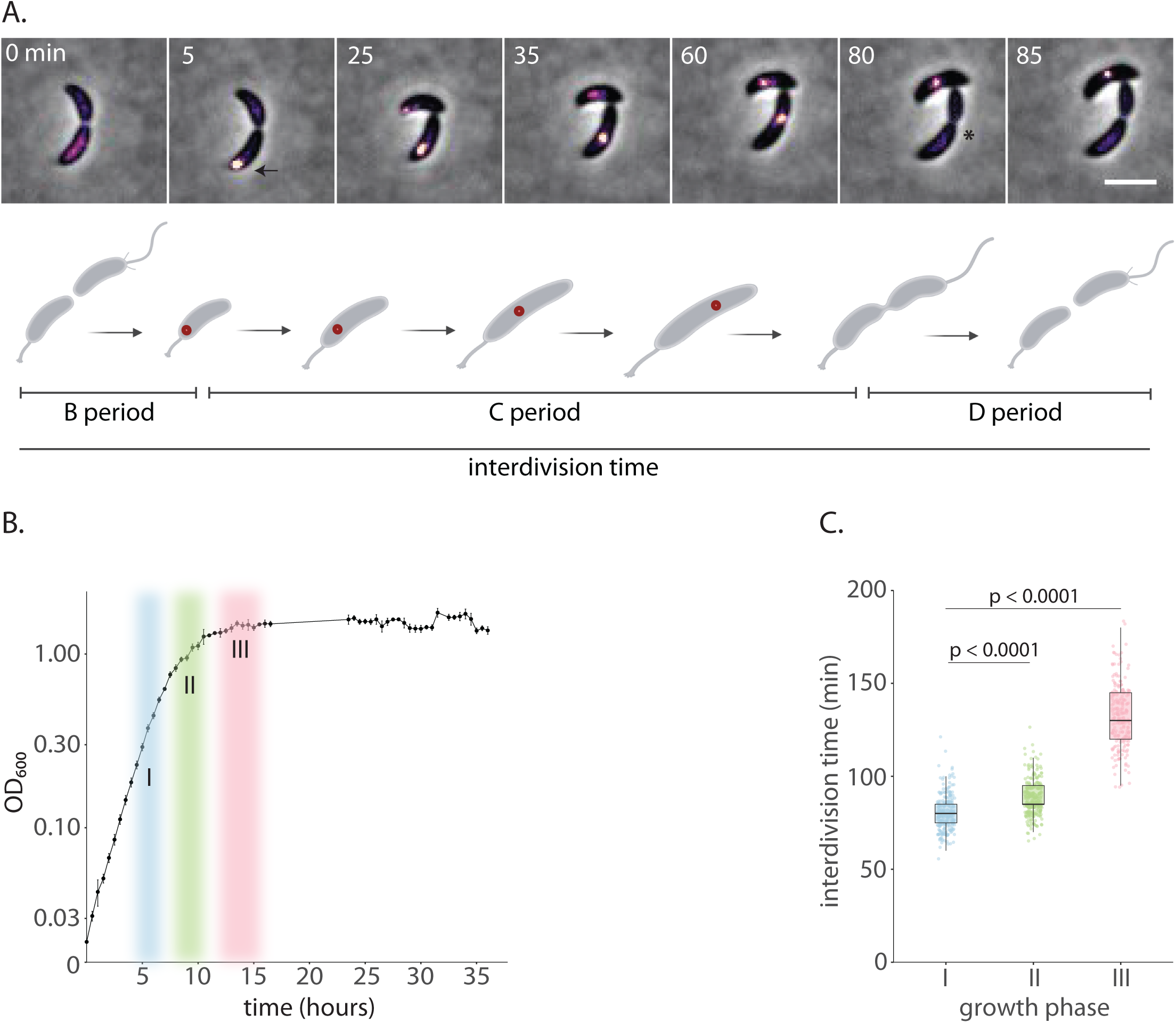
Measurement of *Caulobacter* cell cycle stages via single-cell tracking of replisome dynamics. (A) (Top panel) Representative montage showing progression of replication during *Caulobacter* cell cycle. The montage captures localization of DnaN-YFP (appearing as a focus) at different points of the cell cycle for a cell growing in exponential phase. Time (mins) from the start of the cell cycle is indicated on top left corner of each frame in the montage. Initiation and completion of replication are denoted with arrow and asterisk respectively (scale bar – 2 µm). (Bottom panel) Schematic showing progression of replication during *Caulobacter* cell cycle. Red dots indicate the representative spatial positioning of replisomes along the duration of cell cycle. Duration of each stage of replication (B, C, and D periods) within one cell cycle are also depicted. (B) Growth curve of *Caulobacter crescentus* batch cultures growing in PYE at 30°C. The graph shows mean with SD for optical density measurements (OD_600_) from three independent experiments. The colors denote the growth phases I - mid-exponential, OD_600_ – 0.2-0.4; II – transition from exponential to stationary phase, OD_600_ 0.8-1.4; III – early stationary phase, OD_600_ - 1.4 for a period of 4 hrs) that were characterized for replication dynamics in this study. (C) Interdivision time of bacteria from the three growth phases indicated in 1B. Individual dots in the scatter plot show data from single cells, and the box plots show median with interquartile ranges for each distribution here and in similar graphs elsewhere. The bold line on some of the box plots denote median overlapping with one of the interquartile ranges here and in similar instances elsewhere. n ζ 292 cells pooled from three independent experiments.

We employed this strategy to track cell cycle timing across three distinct phases across the growth curve, with active replication (as deduced by the presence of DnaN localizations) and representing progressively reducing nutrient availability: Phase I - mid-exponential (OD_600_ – 0.2-0.4), Phase II – transition from exponential to stationary phase (OD_600_ 0.8-1.4), and Phase III – early stationary phase (OD_600_ - 1.4 for a period of 4 hrs) (Fig. 1B, S1A-C and S2A). Measurement of interdivision times for these phases indicated that cells slowed down their cell cycle significantly upon entry to stationary phase (interdivision time of 132.4 ± 16.2 min vs 80.2 ± 8.8 min in early stationary and exponential phases respectively) (Fig. 1C, S1A-C).

### Entry into stationary phase is associated with slowdown in replication elongation rates

What factors contribute to this slowdown in cell cycle timing? Previous studies have suggested that *Caulobacter* cells shutdown replication initiation in stationary phase via regulating DnaA levels (Leslie et al., 2015). However, these experiments were conducted much later in stationary phase (approx. 24 hrs after reaching OD_600_ – 1.4), and it is unclear whether initiation shutdown alone can explain an overall slowdown in the cell cycle (as cells still commit to replication at OD_600_ – 1.4). Indeed, two observations from our data suggest an additional layer of replication regulation: 1. after entry into stationary phase (OD_600_ – 1.4), for a period of 6 hrs (∼4 generation times of exponential phase growth), ∼40% of the population harbored replisome foci, and 2. complete replication shut-down (when <5% of the population harbored replisome foci) occurred only much later in stationary phase (24 hrs after reaching OD_600_ – 1.4) (Fig. S2A).

We thus assessed the timing of each replication stage across the three growth phases, to assess the contribution of the same towards the cell cycle time. We found that the B, C and D periods were comparable for Phase I and Phase II cells – B period was found to be 4.6 ± 4 and 6.7 ± 4.1 min, C period 69.8 ± 7.4 and 76.1 ± 7.1 min and, D period 5.8 ± 4.6 and 5.5 ± 4.3 min respectively for Phase I and II (Fig. 2A-C). This is consistent with the observation that interdivision times also did not vary much between these two phases of growth (Fig. 1C). In contrast, upon entry into stationary phase (Phase III), cells significantly slowed down all three replication stages with a B, C and D period of 17.7 ± 9, 103.7 ± 12.9 and 11 ± 6 min respectively (Fig. 2A-C).

**Figure 2:**
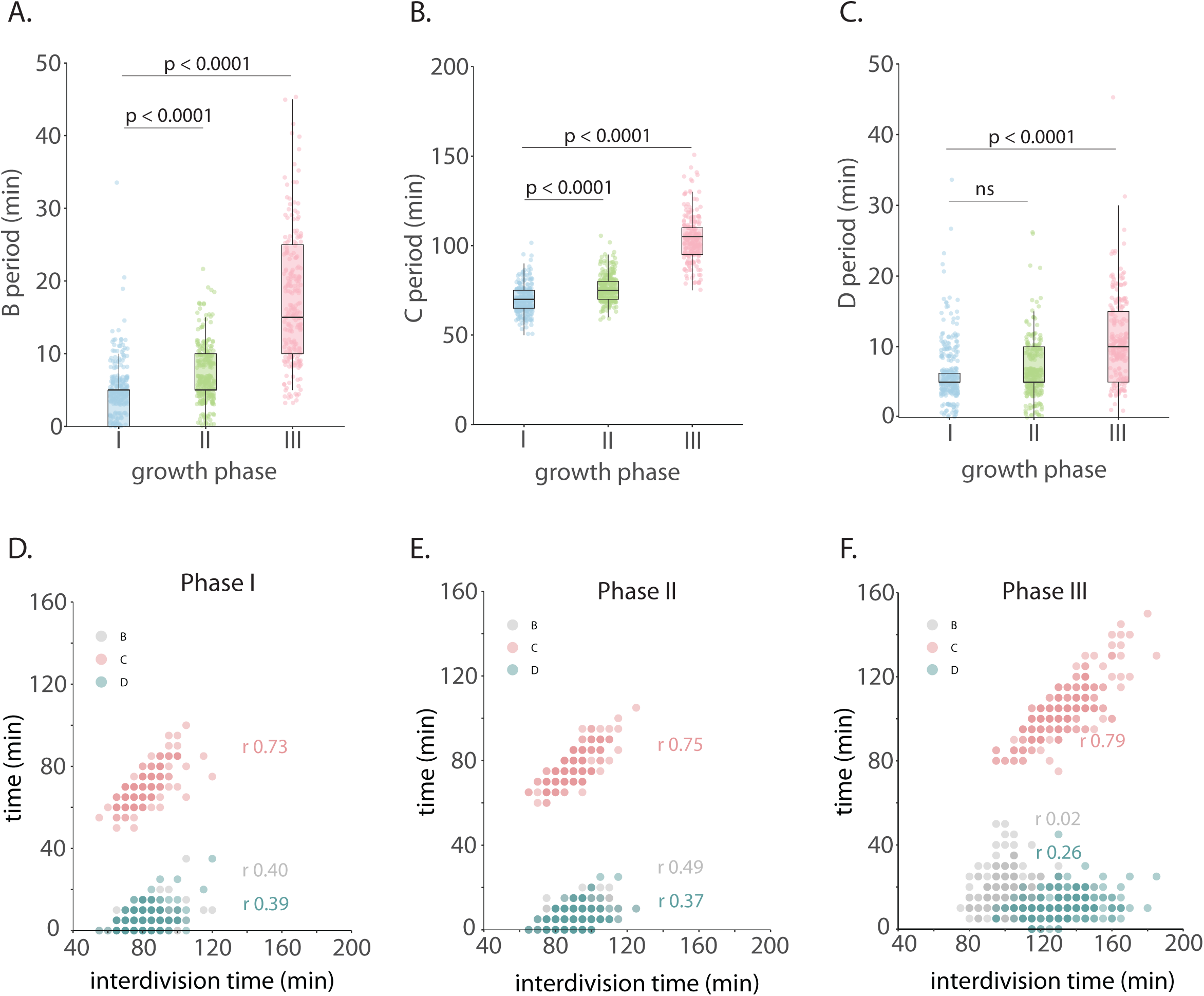
Entry into stationary phase is associated with slowdown in replication elongation rates. (A-C) B, C and D periods of bacteria from the three growth phases indicated in 1B. (D - F) B, C, and D periods of individual bacteria plotted as a function of interdivision time across growth phases I, II, III. n ζ 292 cells pooled from three independent experiments for graphs in A-F. Pearson’s correlation coefficients (r) with CI for B (0.40 [0.30 0.50] for Phase I, 0.49 [0.40 0.57] for Phase II, 0.02 [-0.11 0.15] for Phase III), C (0.73 [0.67 0.78] for Phase I, 0.75 [0.70 0.79] for Phase II, 0.79 [0.73 0.83] for Phase III) and D (0.39 [0.29 0.48] for Phase I, 0.37 [0.27 0.46] for Phase II, 0.26 [0.13 0.37] for Phase III) periods against interdivision time in each growth phase are denoted in respective colors.

In Phase I and II, we observed a weak correlation between interdivision time and B/D periods, though this relationship seemed to break down in Phase III. On the other hand, across all growth phases, the strongest relationship was observed between interdivision time and C period (Pearson’s correlation coefficient (r) 0.73 with confidence interval CI [0.67 0.78], r 0.75 [0.69 0.79] and r 0.79 [0.73 0.83] respectively in growth phases I, II and III) demonstrating the substantial contribution of replication elongation time towards the overall cell cycle duration (Fig. 2D-F).

### Replication slowdown is associated with slower growth rates

Despite the variations in the interdivision/ cell cycle timings, the average length of cells at division were comparable across the three growth phases, with only a modest increase in cell length in growth phases II and III in comparison to growth phase I (mean cell lengths at division being 4.04 ± 0.41 μm, 4.31 ± 0.39 μm and 4.3 ± 0.37 μm for growth phases I, II, and III respectively) (Fig. 4B). We thus asked whether growth rates also slowed down as replication elongation rates increased. For this, we estimated single-cell growth rates (per cell division) for all three growth phases (Fig. 3A). We found that growth rates did reduce from 0.023 ± 0.004 μm/min in exponential phase to 0.015 ± 0.002 μm/min in stationary phase cells (Fig. 3B).

**Figure 3:**
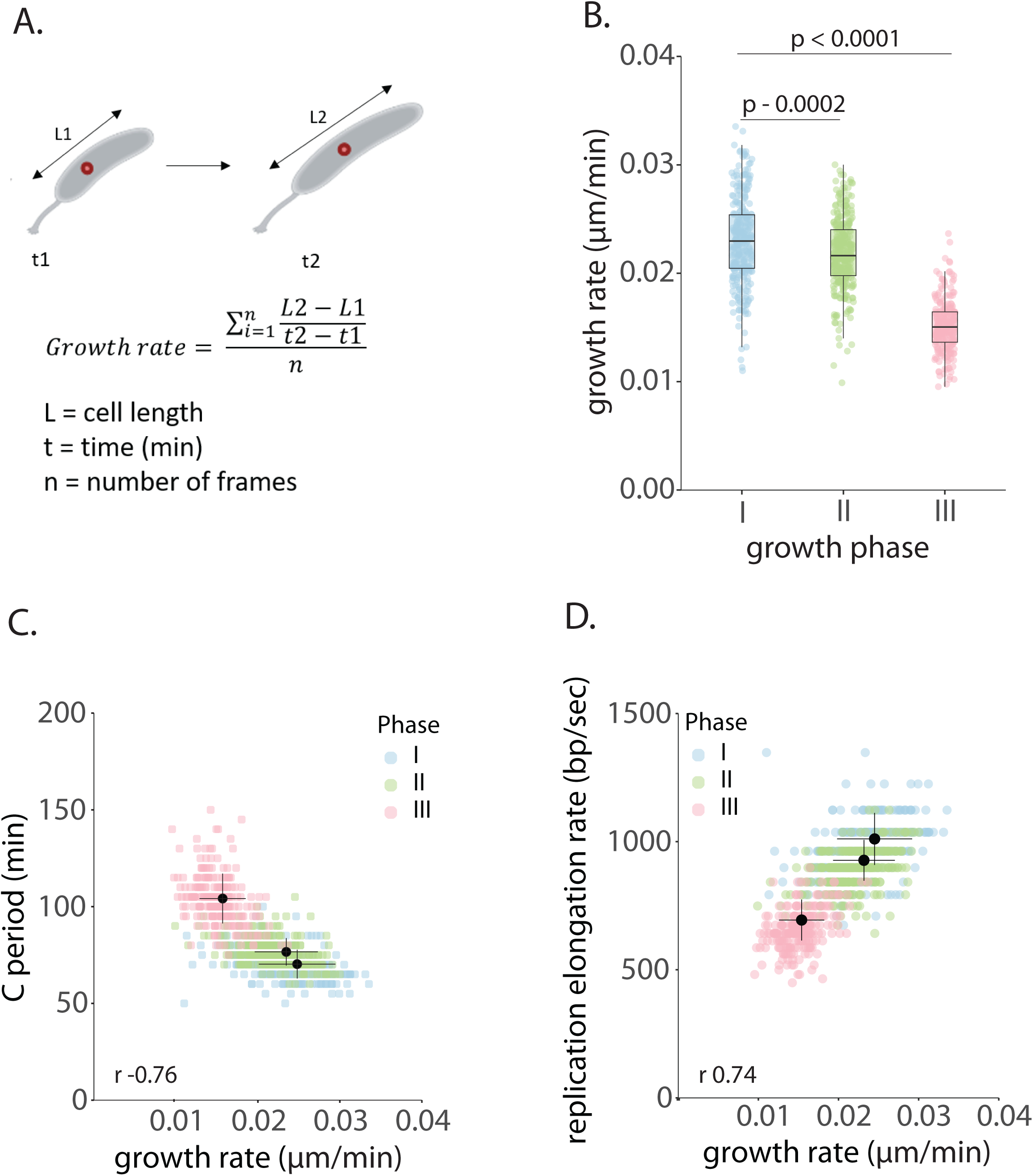
Replication slowdown is associated with slower growth rates. (A) Schematic showing growth rate calculation from single cells. (B) Growth rate of single cells from different growth phases. (C) C periods of single cells from growth phases I, II, III plotted as a function of growth rate. Pearson’s correlation coefficient (r) with CI for C period against growth rate (− 0.76 [-0.77 -0.71]) is indicated in black. (D) Replication elongation rates of single cells from growth phases I, II, III plotted as a function of growth rate. Pearson’s correlation coefficient (r) with CI for replication elongation rate against growth rate (0.74 [0.70 0.77]) is indicated in black. n ý 292 cells pooled from three independent experiments for graphs in A-D.

**Figure 4:**
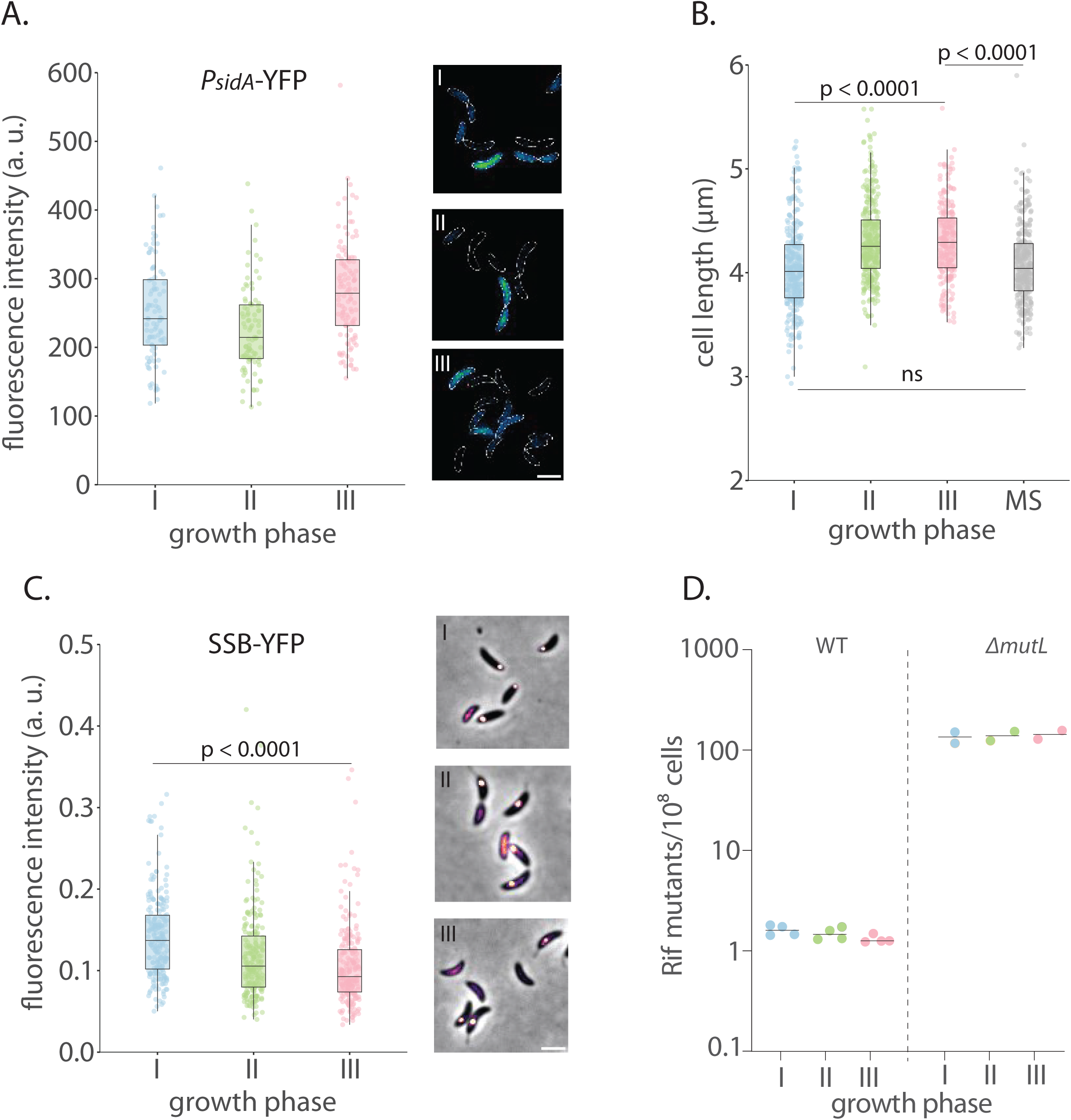
Replication slowdown is not associated with induction of the DNA damage response or increased mutagenesis. (A) (Left) Expression of *yfp* from an SOS-inducible promoter (P*_sidA_*). Total fluorescence intensity normalized to cell area is plotted for bacteria from growth phases I – III. n ý 100 cells pooled from two independent experiments. Fold change in mean fluorescence intensity between phases I and II is 0.9 and that between phases I and III is 1.1 (Right) Representative images of YFP expressed from *P_sidA_* promoter in cells from growth phases I-III (scale bar – 2 µm) (B) Pre-divisional cell lengths of bacteria from growth phases I –III and post-media shift (associated with Fig. S4 and Fig. 6). n ý 292 cells pooled from three independent experiments. (C) (Left) Intensity of SSB-YFP foci normalized to total fluorescence intensity within the cell for bacteria in growth phases I – III. n ý 215 cells from a single experiment. (Right) Representative images of SSB-YFP localization in cells from growth phases I-III (scale bar – 2 µm) (D) Frequency of rifampicin resistant colonies in wild type (WT) and *ΔmutL* cultures from growth phases I - III. Each dot represents frequency calculated from an independent experiment and the black lines indicate medians.

We next compared the cell growth rates with their respective C periods across the three growth phases. We observed a striking relationship between growth rates and replication elongation rates across these populations (r 0.74 [0.70 0.77]), with faster growing cells having a faster replication elongation rate (Fig. 3C-D). We did note that B and D periods additionally showed a weak correlation with growth rate. Cells with slower growth rate also tended to take longer to initiate new rounds of replication or divide after the completion of replication. However, we did not observe any of the individual periods co-varying with each other, suggesting that they are likely regulated independently and/ or by distinct mechanisms (Table S3).

### Replication slowdown is not associated with induction of the DNA damage response or increased mutagenesis

We wondered what factors could contribute to this slowdown in replication elongation rates. Late stationary phase growth has been previously associated with oxidative stress, which can also result in DNA damaging lesions and increased mutagenesis (Gómez-Marroquín et al., 2015; Saumaa et al., 2007). Thus, we considered the possibility of DNA damage as a factor that could perturb replication progression. However, three pieces of evidence suggest that the replisome slowdown observed in our conditions was not associated with DNA damage stress: 1. Cell division was not blocked and cell filamentation was not observed in any of the growth phases (Fig. 4B). In support, cell lengths remained similar across the growth phases as opposed to cell filamentation that occurs in response to DNA damage (Mo & Burkholder, 2010; Modell et al., 2011; Raghunathan et al., 2020) (Fig. 4B). 2. Replication-blocking lesions would reveal long stretches single-stranded DNA (ssDNA) due to continued unwinding of double-stranded DNA by the helicase despite replisome stalling (Thrall et al., 2022) Our measurement of Single-stranded DNA binding protein (SSB) localization intensities did not show a significant increase in cells where replication elongation rate decrease was observed (Fig. 4C). On the contrary, we observed a modest decrease in SSB localization intensities in growth phase III, suggesting a concomitant slowing down of the entire replisome (including the helicase). 3. Substantial SOS response induction was not observed, via assessment of *yfp* expression from the DNA damage-responsive promoter P*_sidA_* (Fig. 4A). In contrast, under DNA damage, the extent of *P_sidA_* induction is much more pronounced (Badrinarayanan et al., 2015; Joseph et al., 2022). Consistently at the onset of stationary phase there was no significant difference in transcript levels of multiple genes that are known to be induced under the SOS response (Table S4).

A second key feature of perturbed replication due to DNA damage is an increase in stress-induced mutagenesis. Indeed, such mutagenesis mediated via error-prone polymerases is reported to occur in the context of late stationary phase cells (grown beyond 48 hrs) (Bull et al., 2001; Sung et al., 2003; Tegova et al., 2004). To assess whether the replication slowdown observed in the conditions tested here is associated with an increase in mutagenesis, we quantified frequency of rifampicin resistant mutants from different phases of growth. We observed similar mutant frequencies across all growth phases tested for wild type cells. Furthermore, ∼100-fold increase in mutation frequencies was observed when mismatch repair was lacking (*ϕ..mutL* cells), and this too remained invariant across all growth phases (Fig. 4D). Together, our observations suggest that DNA damage is an unlikely cause for the observed replication slowdown. Indeed, these data also indicate that replication slowdown itself is not triggering a stress response via SOS induction or increased mutagenesis. Hence, replication elongation rates are affected by the growth environment in a manner that is distinct from replication slowdown under conditions of stress.

### Impact of nutrient status on replication progression

Another factor that could contribute to the observed slowdown in replication elongation rates could be nutrient availability. Transition from exponential to stationary phase is characterized by alterations in cellular physiology due to shifts in nutrient availability and metabolism, which can potentially impact growth dynamics of bacteria (Bergkessel & Delavaine, 2021; Hengge, 2011; Navarro Llorens et al., 2010). We thus asked whether variation in nutrient status in the growth medium would reflect in variation in replication elongation rates. For this, we compared replication progression in exponentially growing steady-state cells in PYE (rich media) vs minimal media with two defined carbon sources (M2G - with 0.2% glucose and M2X - with 0.2% xylose) (S3A).

We observed that cells grown in M2G or M2X had significantly longer C periods when compared to PYE, with M2X cells showing the slowest replication elongation rates (976 ± 101, 751 ± 75 and 722 ± 73 bp/sec for PYE, M2G and M2X respectively) (Fig. 5A-D). Consistently, measurement of interdivision time revealed longer cell cycle duration in minimal media conditions (Fig. 5E) and the correlation between C period and growth rates were also recapitulated (r -0.78 [-0.81 -0.76]) (Fig. 5C). As in the case of different growth phases, interdivision time displayed strongest relationship with C period compared to B or D periods (Fig. 2D-F, S3B-C). These data are indicative of a link between nutrient availability, growth rates and replication elongation rates.

**Figure 5:**
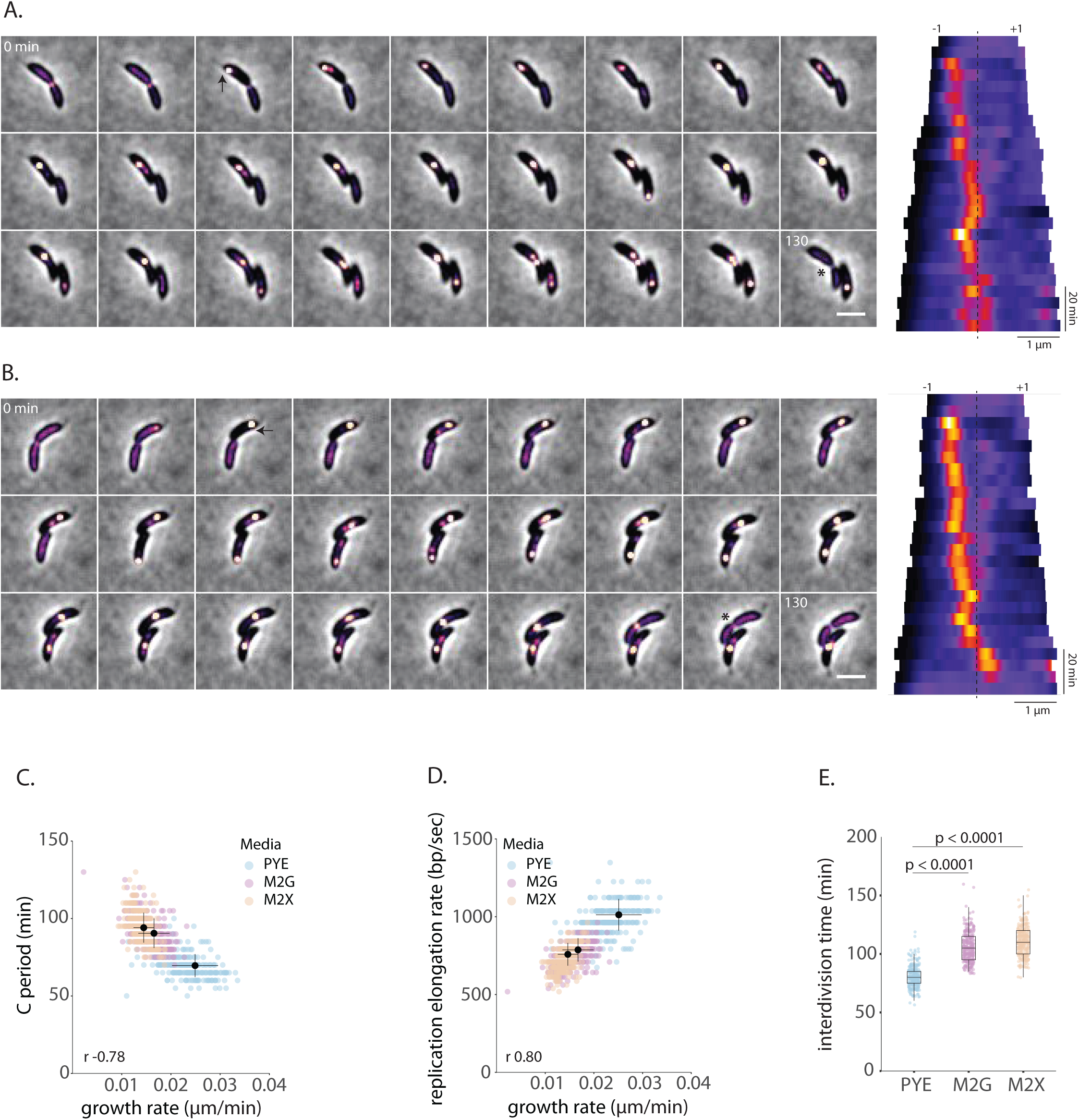
Impact of nutrient status on replication progression. (A-B) Representative time-lapse montages and corresponding kymographs showing replication progression (DnaN-YFP localization) in cells growing in M2G and M2X. Time (mins) from the start of the cell cycle is indicated on top left corner of the first and last frame of each montage. Initiation and completion of replication are denoted with arrows and asterisks respectively in the montage (imaging interval – 5 min, scale bar – 2 µm). In kymographs mid cell position is indicated with the dashed line. Old and new poles are denoted with -1 and +1 respectively. (C) C periods of single cells growing in PYE, M2G or M2X plotted as a function of growth rate. Pearson’s correlation coefficient (r) with CI for C period against growth rate (−0.78 [-0.81 -0.76]) is indicated in black. (D) Replication elongation rates of single cells growing in PYE, M2G or M2X plotted as a function of growth rate. Pearson’s correlation coefficient (r) with CI for replication elongation rate against growth rate (0.80 [0.78 0.83]) is indicated in black. Data for PYE phase I in C and D has been reproduced from 3C and 3D for comparison. (E) Interdivision time of bacteria growing in PYE, M2G and M2X. Data for PYE (Phase I) has been reproduced from Fig. 1C for comparison. n ≥ 292 cells pooled from three independent experiments for graphs in 3C-E.

### Replication elongation rates robustly change with alteration in nutrient availability

To directly assess the impact of nutrient availability on replication elongation rates, we designed an experiment to supplement stationary phase cells with additional nutrients in the growth media. For this, we shifted cells growing in stationary phase (OD_600_ – 1.4) into growth media extracted from exponentially growing cells (OD_600_ − 0.2), and assessed the timing of the very first cell cycle of these ‘media-shifted’ cells in their new growth environment (Fig. S4A). We observed an immediate reduction in the C period of these media-shifted cells (103.7 ± 12.9 min and 85.3 ± 12 min in stationary and media-shifted cells respectively). This was also reflected in the interdivision time of these media-shifted cells from 132.4 ± 16.2 min in stationary phase to 102.6 ± 16.2 min in media-shifted conditions (Fig. 6A-B). We did also observe an increase in growth rates, but not to the same extent as seen for the change in interdivision time or C period (growth rate of 0.015 ± 0.002 and 0.017 ± 0.004 μm/min in stationary phase and media-shifted cells respectively) (Fig. 6C-D). Indeed, this decoupling was also reflected in the cell lengths, with media-shifted cells having a modestly smaller mean cell length when compared with stationary phase cells (4.08 ± 0.37 and 4.3 ± 0.37 μm respectively) (Fig. 4B). One possibility is that nutrient availability independently influences replication and growth rates. This is also consistent with observations in cells transitioning from exponential phase towards stationary phase. In this case we found that the proportion of cells initiating replication already started to decline from mid-exponential phase even though growth rates remained unchanged until the transition from exponential to stationary phase (Fig. S2A). Together, our observations are consistent with replication speed variation in a nutrient-dependent manner.

**Figure 6:**
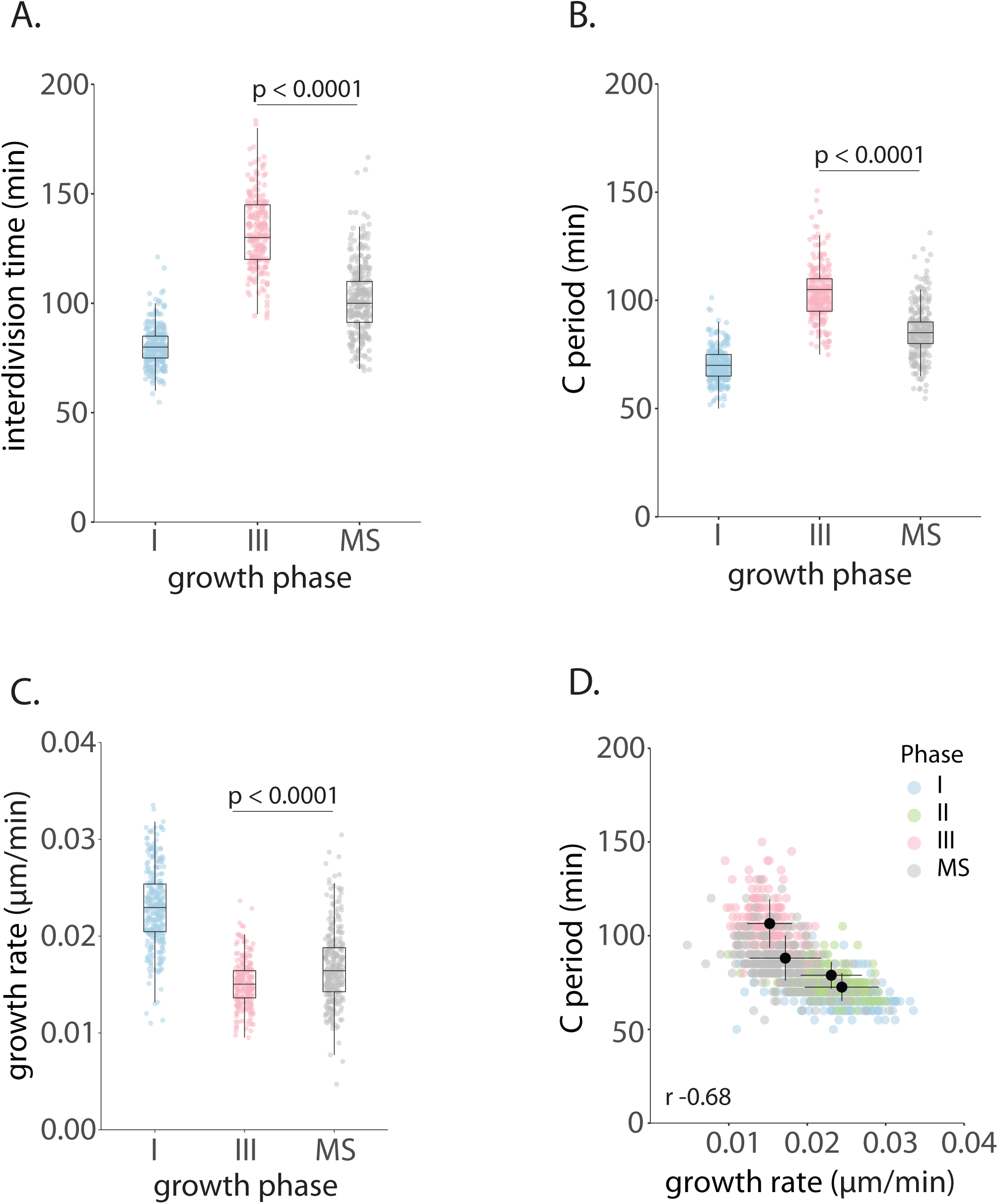
Replication elongation rates robustly change with alteration in nutrient availability. (A) Interdivision time of bacteria from growth phases I and III compared to those shifted from growth phase III to media from growth phase I (MS – media shift). Data for growth phases I and III has been reproduced from Fig. 1C for comparison. (B) C periods of bacteria from growth phases I and III compared to those shifted from growth phase III to media from growth phase I (MS). Data for growth phases I and III has been reproduced from Fig. 2B for comparison. (C) Growth rate of single cells from growth phases I and III compared to those shifted from growth phase III to media from growth phase I (MS). Data for growth phases I and III has been reproduced from Fig. 3B for comparison. (D) C periods of bacteria shifted from growth phase III to media from growth phase I (MS) plotted as a function of growth rate. Data for growth phases I – III has been reproduced from Fig. 3C for comparison. Pearson’s correlation coefficient (r) with CI for C period against growth rate (−0.68 [-0.71 -0.65]) is indicated in black.

## Discussion

Response of individual bacteria to fluctuations in growth environment (such as nutrient availability) can collectively shape the fitness of bacterial populations and diversity within microbial communities (Fenchel, 2002; Nguyen et al., 2021). The most frequent responses to many environmental fluctuations include alterations in bacterial metabolism, growth and cell cycle; whereas resource abundance results in fast growth and division, resource depletion leads to slowing down or inhibition of both (Bergkessel & Delavaine, 2021; Nguyen et al., 2021; Wang & Levin, 2009).

The transition between fast and slow growth modes as observed within natural niches can be recapitulated to some extent in batch cultures of bacteria growing under optimal laboratory conditions. As resource depletion and other stresses set in, the exponentially multiplying bacterial population enters stationary phase, where bacterial physiology changes, and the viable bacterial count in the population starts to stagnate (Hengge, 2011; Navarro Llorens et al., 2010; Nyström, 2004). In fact, in nature bacteria are thought to go through similar cycles of feast and famine, with a large proportion of bacteria being in physiological states comparable to stationary phase (Hengge, 2011; Kolter et al., 1993; Nyström, 2004). The progressive deterioration in growth conditions during exponential to stationary phase transition provides an excellent window to assess temporal regulation of cellular processes during transitions between growth modes.

Indeed, during transitions between growth modes, one of the most critical processes to be regulated is DNA replication. To ensure a complete duplicated copy of the genome prior to every division, it is imperative that replication is coordinated with growth and cell division. In our current work, we systematically characterized replication dynamics in single *Caulobacter* cells, across different growth contexts encompassing bacterial growth phases as well as media conditions. Our findings directly demonstrate at the level of single cells, the regulation of replication elongation rates on transition from exponential to stationary phase, which is also recapitulated under media conditions of varied richness. This work has revealed a clear link between nutrient status, bacterial growth and rate of genome duplication. We consider a few possible mechanisms driving this regulation of replication elongation rates below.

### Nutrient-dependent regulation of replication

Although the speed of DNA synthesis is inherent to the polymerase, multiple factors can affect the processivity of the holoenzyme, including stoichiometry of essential subunits of the replisome and status of dNTP precursors. For instance, earlier studies in *E. coli* have demonstrated that titrating the expression of ribonucleotide reductase (RNR) enzymes or inhibiting this enzyme with hydroxyurea can lead to alterations in dNTP pools and substantial reduction in the C period duration (Odsbu et al., 2009; Zhu et al., 2017). However, we did not observe significant differences in transcript levels of RNR genes involved in dNTP biosynthesis (*nrdE, nrdB, nrdJ or CCNA_01420*) between bacteria in exponential phase (Phase I) and the onset of stationary phase (Phase III) (Table S4).

Another consideration is the role of the alarmone (p)ppGpp which mediates stringent response in bacteria. It has been observed that induction of the stringent response affects not only replication initiation (Fernández-Coll et al., 2020; Gonzalez & Collier, 2014; Gross & Konieczny, 2020; Kraemer et al., 2019; Lesley & Shapiro, 2008), but also elongation rates. (DeNapoli et al., 2013; Giramma et al., 2021; Wang et al., 2007). In *Bacillus subtilis* and *E. coli* (p)ppGpp inhibits the activity of primase DnaG, either by binding to it or lowering levels of GTP. However, (p)ppGpp production and associated stringent response are generally triggered under severe carbon and amino acid deprivation (DeNapoli et al., 2013; Giramma et al., 2021; Lesley & Shapiro, 2008; Wang et al., 2007). It is unlikely that the growth phases and minimal media conditions used in the current study would result in such drastic nutrient deprivation, and hence the differences observed in replication time might not be mediated by (p)ppGpp. Moreover, in *Caulobacter*, the deletion of *spoT*, responsible for (p)ppGpp production does not impact replication initiation even at a later stage of stationary phase than the one studied here (Leslie et al., 2015). Our own RNA-seq experiments further indicate that the expression of (p)ppGpp synthase gene (*spoT/CCNA_01622*) is not induced at the onset of stationary phase (Phase III) (Table S4). However, expression levels of *spoT* alone may not be indicative of cellular (p)ppGpp levels. Hence, further investigations are essential to clarify if the physiological growth contexts studied here have significantly impacted dNTP pool and/ or (p)ppGpp activity that could result in an altered C period in *Caulobacter*.

Our data indicate that even though replication slows down at the onset of stationary phase, the ssDNA in front of the replication forks does not increase. Indeed, the modest reduction in SSB foci intensities in stationary phase would suggest a concomitant slowdown of the helicase as well. It is possible that replication elongation rates in different growth phases could be regulated by the rate of DNA unwinding by the helicase whose function is ATP dependent.

### Going fast and slow

Variations in replication rates can have important consequences. On the one hand, given the trade-off between speed and accuracy, it can be anticipated that slow replisomes may have lower mutation rates (Banerjee et al., 2017; Bhat et al., 2022; Niccum et al., 2019). On the other hand, slow replisomes can result in generation of longer stretches of ssDNA between the stalled replisome and the helicase unwinding the double-stranded DNA ahead of it. This could trigger induction of the SOS response, that results in the expression of error-prone polymerases contributing to an increase in mutagenesis. For example, treatment of cells with hydroxyurea which depletes cellular dNTP pools can lead to generation of ssDNA tracts that elicit an SOS response (Thrall et al., 2022).

Instead, our observations suggest that there is a ‘Goldilocks’ zone of replication rates with large range (660 ± 80 to 976 ± 101 bp/sec in present study) in which variation is tolerated without significant impact on mutagenesis. Furthermore, these rates are also readily reversible (dependent on nutrient availability) indicating the plasticity of this process. It is possible that there are tolerable levels of environmental fluctuations within which such plasticity is observed and beyond a threshold, fluctuations could be associated with more permanent changes. Thus, while replication slowdown mediated by genotoxic stress is often associated with activation of stress responses and stress-induced mutagenesis (Tan et al., 2015; Thrall et al., 2022), replisome speed variation that occurs in response to nutrient availability (such as observed here) is likely a robust adaptation to fluctuating environmental conditions,

We speculate that such changes in replication elongation rates might be essential to coordinate replication with cell growth as well as other processes that rely on metabolic states. This could also contribute to a prolonged stationary phase (observed here) where replication is still active, prior to a complete shutdown. Such slowdown could enable cells to rapidly return to fast growth in case nutrients become readily available in the environment. In human cell line studies, it was observed that the inability of replication forks to sense cellular metabolic states and slowdown replication resulted in loss of genome integrity, highlighting the importance of modulating replication elongation rates (Somyajit et al., 2017). Similarly in bacteria, nutrient-dependent regulation of replication rates could be a strategy to cope with transient fluctuations in natural niches without requiring to induce stress responses or mutagenesis. The mechanisms that regulate replication rates and/ or the threshold of nutrient fluctuations beyond which stress responses and mutagenesis are induced are exciting and important open questions.

## Materials and Methods

### Bacterial strains and growth conditions

Details of bacterial strains and plasmids used in this study are available in Supplementary file (Tables S1 & S2). *Caulobacter crescentus* strains were routinely grown in PYE (0.2% peptone, 0.1% yeast extract and 0.06% MgSO4) supplemented with antibiotics as required. While growing strain expressing SSB-YFP under *P_xyl_*, 0.3% xylose was added to the culture 3 hrs before the experiment commenced. For experiments in minimal media bacteria were grown in M2 minimal media (1X M2 salts - 0.087% Na2HPO4, 0.53% KH2PO4, and 0.05% NH4Cl, 0.01 mM FeSO4 and 0.01 mM CaCl2) supplemented with either 0.2% glucose (M2G) or 0.2% xylose (M2X).

### Growth curve measurements

For growth curve measurements, overnight cultures were back diluted to OD_600_ – 0.025 (PYE) or 0.05 (M2G and M2X). OD_600_ measurements were recorded manually every half an hour using Hitachi UH5300 Spectrophotometer. The measurements were carried out in at least three independent cultures under each growth/media regime. For growth curve, mean with SD from three independent experiments were plotted.

### Fluorescence microscopy – sample preparation and imaging

For time course microscopy, aliquots were taken from cultures growing at specified conditions at different time points, pelleted down and resuspended in appropriate volumes of growth medium. 2 μl of the cell suspension was spotted on 1% agarose pads (prepared in water) with approximate dimensions of 0.5 X 0.5 cm.

For time lapse microscopy, two independent cultures were grown to the same optical density – one for imaging and the other for extracting spent media. The culture used for extraction of spent media was grown to required OD_600_ approximately 3 hrs ahead of the culture used for imaging. This culture was then filtered with a 0.22 µm membrane filter and the filtrate (spent media) was used to prepare 1.5% GTG agarose pads. At specified OD_600_ an aliquot taken from the culture to be imaged was pelleted down, washed and resuspended in the spent media. 2 μl of the cell suspension was spotted on the pads (of similar dimensions as mentioned above), grown inside an OkoLab incubation chamber maintained at 30°C and imaged at 5 min intervals for a duration of 6 hrs.

Imaging was done on Nikon Eclipse Ti2 wide-field microscope equipped with a motorized XY stage, using a 60X plan apochromat objective (NA 1.4) and illumination from pE4000 light source (CoolLED). Image capture was performed with Hamamatsu Orca Flash 4 camera using Nikon’s NIS Elements software. Infrared-based Perfect Focusing System ensured maintenance of focus during time lapse imaging. Fluorescence imaging of YFP and mCherry were done with excitation at 490 nm and 550 nm respectively, for exposure times of 400 ms.

### Image analysis

Percentage cells with DnaN foci from time course microscopy was manually analysed using CellCounter plugin on ImageJ. For calculating interdivision time, B, C and D periods from time lapse imaging, independent crops of cells in which complete cell cycle could be tracked were made. These crops were manually analysed for each period and interdivision time using ImageJ. For cell length analysis, the same cell crops were segmented using Oufti, and cell lengths were extracted using a MATLAB (version – 2020a) script. The same data was used to analyse growth rate using the formula indicated in Fig. 3A. Replication rates were calculated by dividing whole genome length by replication time. For fluorescence intensity analysis of YFP expression from *P_sidA_* promoter, cells were segmented with MicrobeJ plugin on ImageJ and total fluorescence intensity normalized to cell area was extracted. For calculating SSB foci intensity, each focus was segmented using MicrobeJ plugin on ImageJ with maxima (foci) function, and total intensity of each focus was normalized to total fluorescence intensity within that cell.

### Statistical analysis

All graphs were plotted on RStudio (R version – 4.3.3). For calculating statistical significance (Fig. 1C, 2A-C, 3B, 4A-C, 6A-C, 5E) data from three independent biological replicates were pooled, and outliers were first removed using the z-score method. Brown-Forsythe test was used to ensure similarity in variance and one-way ANOVA with Bonferroni correction was performed (assuming these distributions to be normal). Pearson’s correlation coefficients (r) were obtained to assess correlations in Fig. 2D-F, 3C-D, 5C-D, S3B-C, S4B. Results of all statistical tests are consolidated in Table S3.

### RNA Sequencing

1.5 ml aliquots of CB15N cultures in exponential (OD_600_ – 0.4) and stationary (OD_600_ – 1.4) phases were collected, pelleted down, snap frozen in liquid N_2_ and stored at -80°C until further processing. RNA was isolated from these pellets using Direct-zol RNA Miniprep kit (Zymo Research) as previously described (Kamat et al., 2024). Post-DNase treatment to remove any genomic DNA, these RNA samples were purified using RNA Clean and Concentrator-25 columns (Zymo Research). In order to enrich mRNA pool in these samples, they were subjected to rRNA depletion using Ribominus rRNA depletion kit and further purified using RNA Clean and Concentrator-5 columns (Zymo Research). RNA integrity was checked using Bioanalyzer before proceeding to library prep and sequencing. Library prep was done with NEBNext® Ultra™ II Directional RNA Library Prep Kit and paired end sequencing was carried out on Illumina Novaseq6000 sequencing platform at NCBS NGS facility. Raw reads obtained as Fastq files were processed using a previously established pipeline to retrieve mapped read counts per gene (Kamat et al., 2024). TMM normalized reads were used to calculate log-fold-changes (Log_2_FC) in stationary phase compared to exponential phase. Log_2_FC >1 was considered upregulated (Table S4).

### Mutagenesis assay

Overnight cultures of WT or */-mutL* strains were back diluted to OD_600_ of 0.1. Aliquots were drawn from these cultures at OD_600_ – 0.4, 0.8 and 1.4, and plated on 100 μg/ml rifampicin plate. At each of these time points a separate aliquot was drawn to calculate viable counts in the volume of culture plated on rifampicin plate. Colonies were counted after 48 hrs of incubation at 30°C. Rif mutant frequencies were calculated by normalizing Rif resistant colonies to viable counts in each culture.

## Author contributions

AJ and AB conceived the project. IA led the project, carried out majority of the experiments, generated tools and conducted data analysis. SP carried out experiments across different growth media and conducted analysis of the same. AJ conducted mutagenesis experiments. AJ and AB supervised the project. AB procured funding. AJ and AB wrote the manuscript, with inputs from all authors.

## Acknowledgements

The authors thank AB lab members for feedback on the work. This work was supported by funding to IA and SP (TIFR graduate student fellowship) and to AB via the India Alliance Intermediate Grant (Grant number IA/I/21/1/505630) and intramural funding via NCBS-TIFR (Department of Atomic Energy, Government of India, Project Identification No. RTI 4006).

## Data availability

All sequencing data are deposited in GEO

---

**Table S1:** Strains used in the study

| Strain name | Genotype | Strain construction |
| --- | --- | --- |
| CB15N | *NA1000* |  |
| NABC124 | *CB15N; dnaN-YFP::spec<sup>R</sup>* | CB15N was transformed with pNABC198 plasmid to integrate *dnaN-YFP* linked to *spec<sup>R</sup>* at endogenous locus |
| NABC12 | *CB15N; dnaN-mCherry::kan<sup>R</sup>* | CB15N strain with *dnaN-mCherry* linked to *kan<sup>R</sup>* integrated at the endogenous locus. |
| NABC632 | *CB15N; Pxyl-ssb-YFP::kan<sup>R</sup>* | CB15N transformed with pNABC419 plasmid to integrate *ssb-YFP* linked to *kan<sup>R</sup>* under the *P<sub>xyl</sub>* promoter at *xyl* locus |
| NABC581 | *CB15N; P<sub>sidA</sub>-YFP::kan<sup>R</sup>* | Joseph *et al.*, 2022 |
| NABC580 | *CB15N; ΔmutL* | Joseph *et al.*, 2022 |

**Table S2:** Plasmids used in the study

| Plasmid name | Construct details | Antibiotic resistance |
| --- | --- | --- |
| pNABC198 | Joseph *et al.*, 2021 | Spectinomycin |
| pNABC419 | Joseph *et al.*, 2021 | Kanamycin |

**Figure S1:**
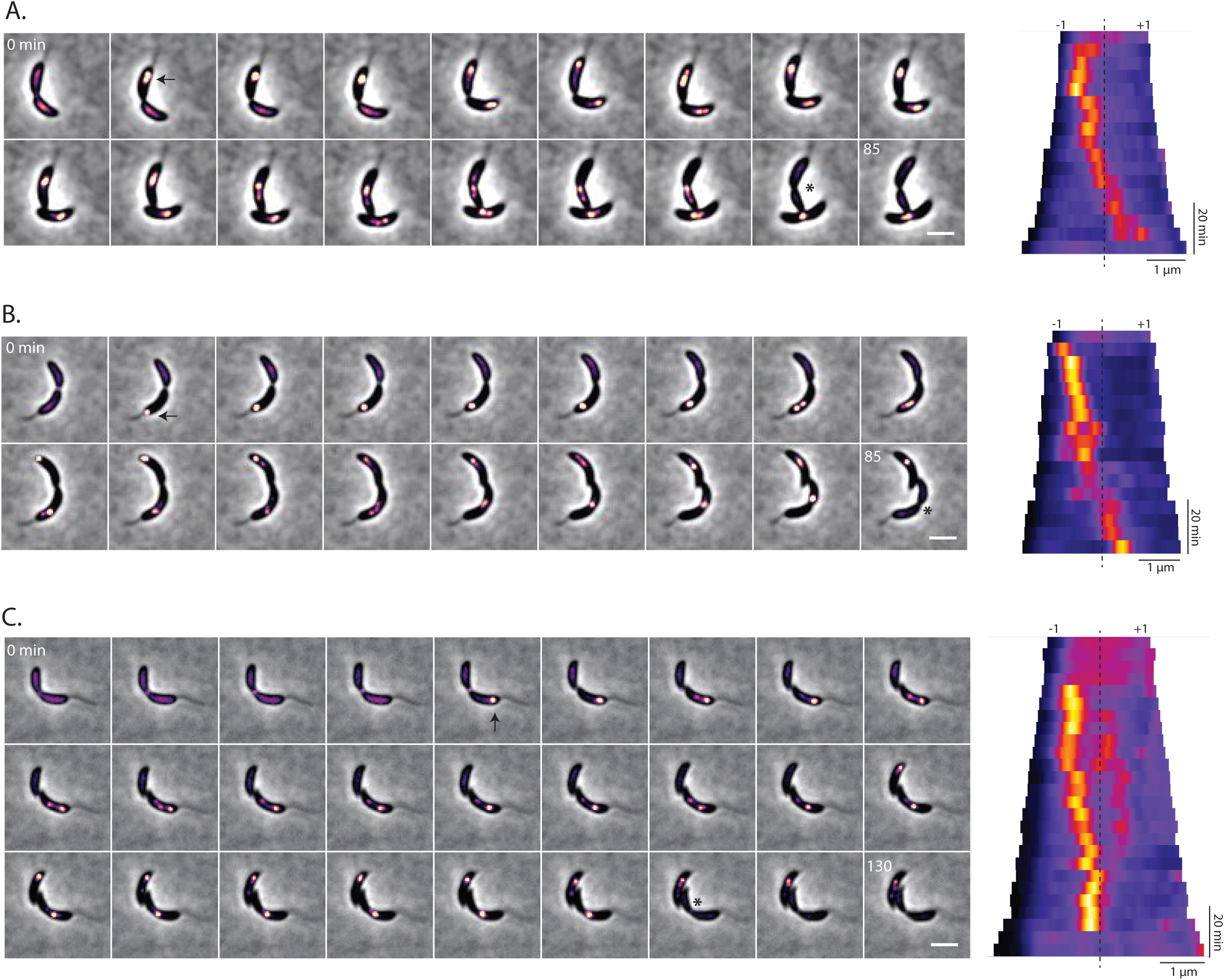
Measurement of *Caulobacter* cell cycle stages via single-cell tracking of replisome dynamics. (A-C) Representative time-lapse montages and corresponding kymographs showing replication progression (DnaN-YFP localization) in cells from growth phases I-III. Time (mins) from the start of the cell cycle is indicated on top left corner of the first and last frame of each montage. Initiation and completion of replication are denoted with arrows and asterisks respectively in the montage (imaging interval – 5 min, scale bar – 2 µm). In kymographs mid-cell position is indicated with the dashed line. Old and new poles are denoted with -1 and +1 respectively.

**Figure S2:**
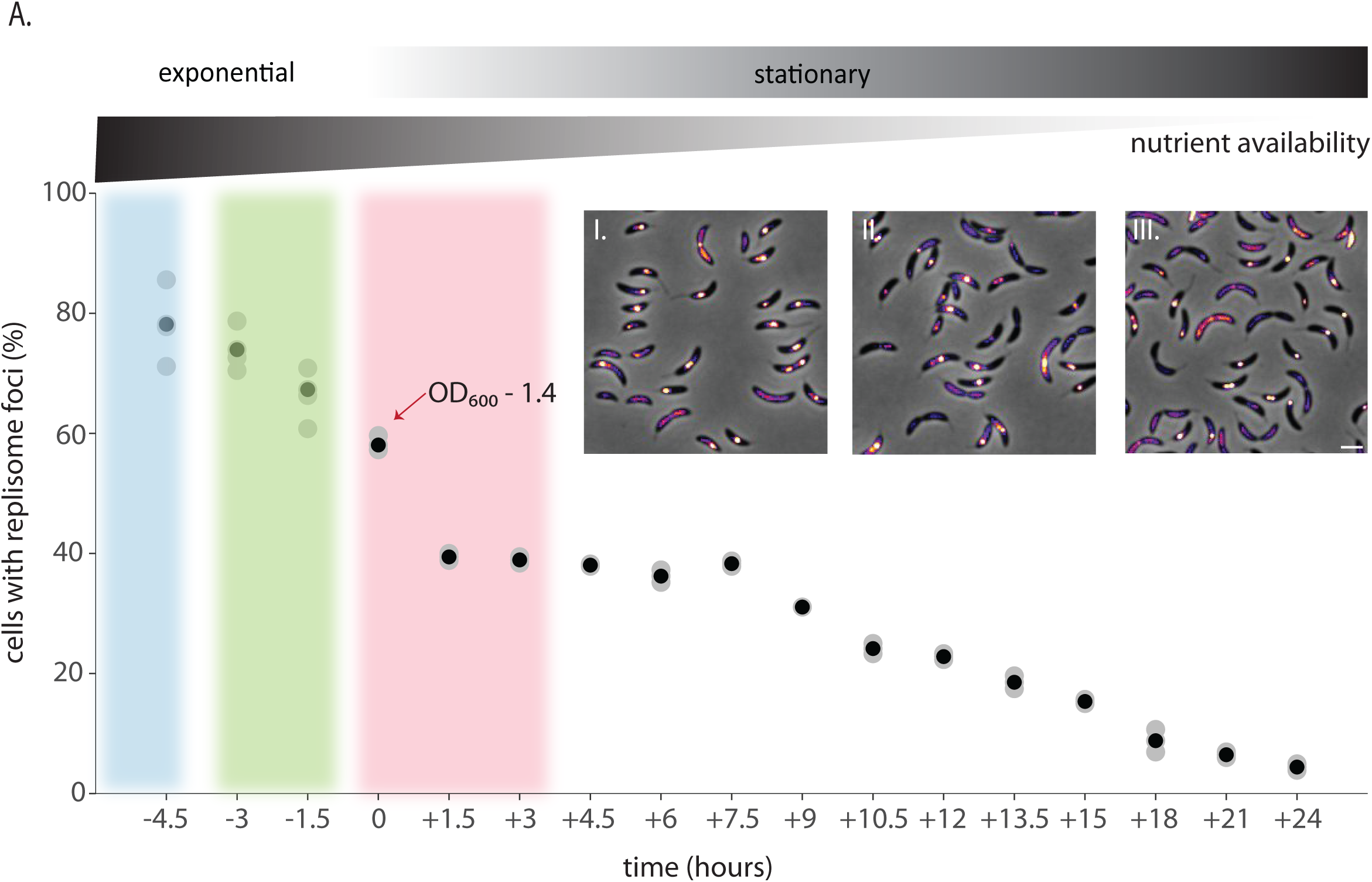
Entry into stationary phase is associated with slowdown in replication elongation rates. (A) Percentage cells harboring DnaN-mCherry foci at different points in the growth curve depicted in 1B. Grey circles denote independent experiments, and the black dots indicate mean for each time point. n ≥ 200 cells pooled from at least two independent experiments. (Inset) Representative snapshots showing localization of DnaN-mCherry in growth phases I-III (scale bar -2 µm).

**Figure S3:**
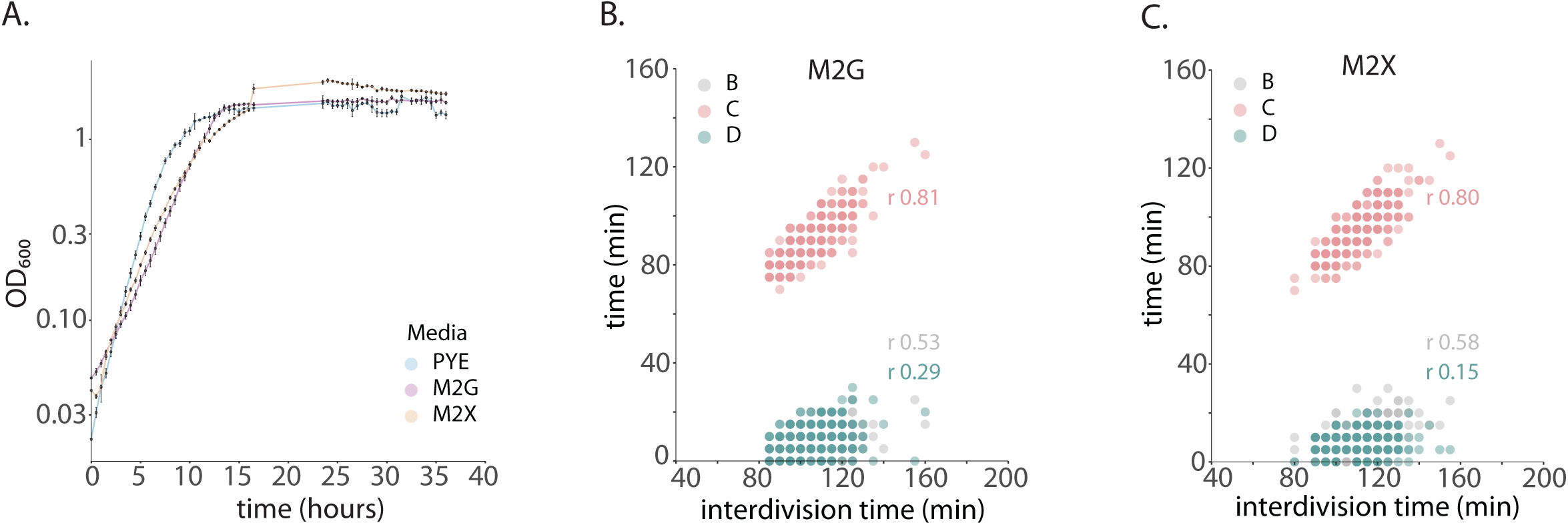
Impact of nutrient status on replication progression. (A) Growth curves of *Caulobacter crescentus* batch cultures growing in PYE, M2G and M2X at 30°C. The graph shows mean with SD of optical density measurements (OD_600_) from three independent experiments. Data for PYE growth curve has been reproduced from Fig. 1B for comparison. (B-C) B, C, and D periods of individual bacteria plotted as a function of interdivision time for bacteria growing in M2G or M2X. n ≥ 300 cells pooled from three independent experiments. Pearson’s correlation coefficients (r) with CI for B (0.53 [0.45 0.610] for M2G, 0.58 [0.50 0.65] for M2X), C (0.81 [0.77 0.85] for M2G, 0.80 [0.76 0.84] for M2X) and D (0.29 [0.18 0.39] for M2G, 0.15 [0.03 0.25] for M2X) periods against interdivision time are denoted in respective colors.

**Figure S4:**
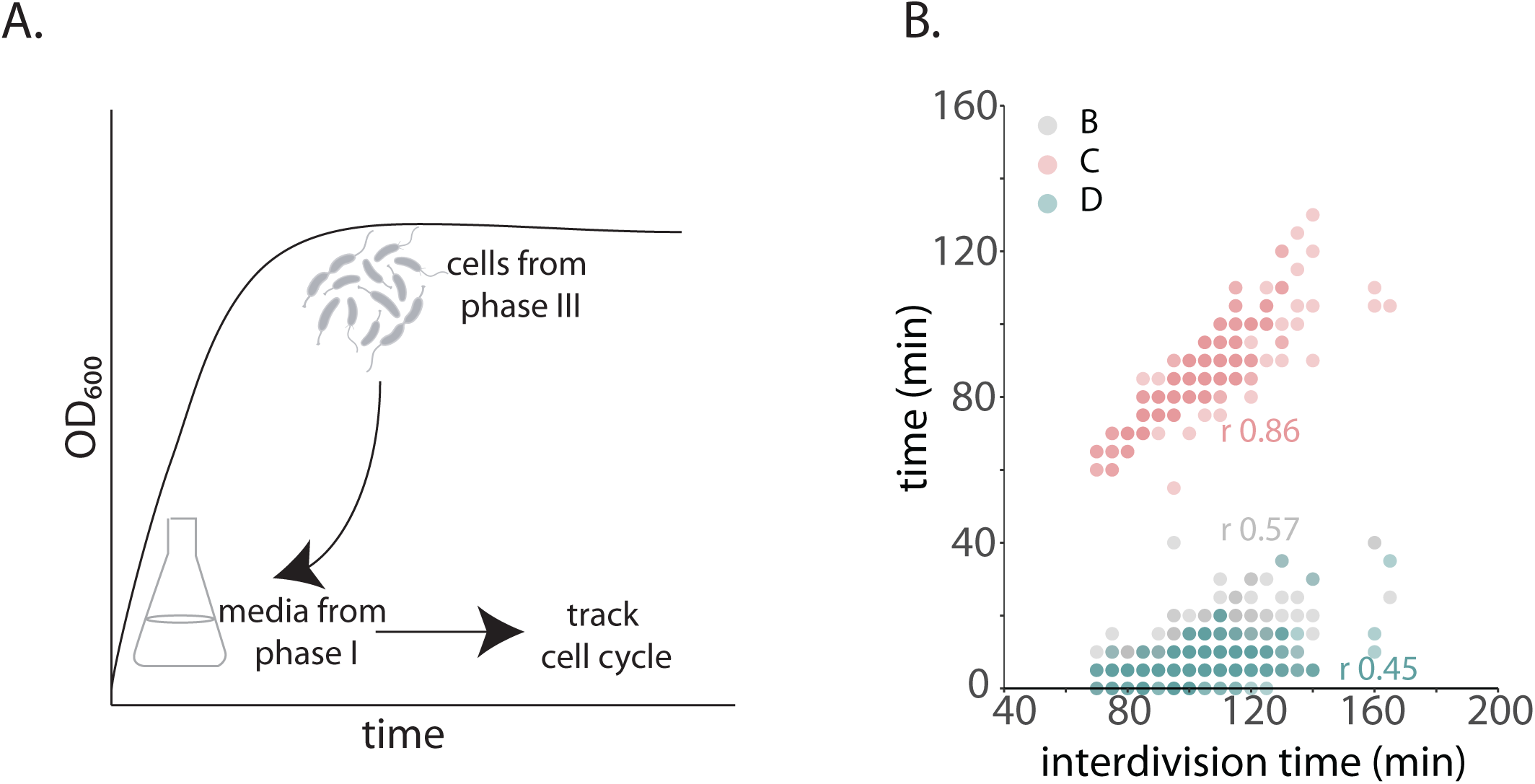
Replication elongation rates robustly change with alteration in nutrient availability. **(A)** Schematic showing experimental set-up for media shift experiment where stationary phase cells are shifted to media extracted from exponential phase. (B) B, C, and D periods of individual bacteria plotted as a function of interdivision time for media-shifted cells. n = 300 cells pooled from three independent experiments. Pearson’s correlation coefficients (r) with CI for B (0.57 [0.49 0.64]), C (0.86 [0.82 0.88]) and D (0.45 [0.36 0.54]) periods against interdivision time are denoted in respective colors.

